# From Noise to Knowledge: Diffusion Probabilistic Model-Based Neural Inference of Gene Regulatory Networks

**DOI:** 10.1101/2023.11.05.565675

**Authors:** Hao Zhu, Donna K. Slonim

## Abstract

Understanding gene regulatory networks (GRNs) is crucial for elucidating cellular mechanisms and advancing therapeutic interventions. Original methods for GRN inference from bulk expression data often struggled with the high dimensionality and inherent noise in the data. Here we introduce RegDiffusion, a new class of Denoising Diffusion Probabilistic Models focusing on the regulatory effects among feature variables. RegDiffusion introduces Gaussian noise to the input data following a diffusion schedule and uses a neural network with a parameterized adjacency matrix to predict the added noise. We show that using this process, GRNs can be learned effectively with a surprisingly simple model architecture. In our benchmark experiments, RegDiffusion shows superior performance compared to several baseline methods in multiple datasets. We also demonstrate that RegDiffusion can infer biologically meaningful regulatory networks from real-world single-cell data sets with over 15,000 genes in under 5 minutes. This work not only introduces a fresh perspective on GRN inference but also highlights the promising capacity of diffusion-based models in the area of single-cell analysis. The RegDiffusion software package and experiment data are available at https://github.com/TuftsBCB/RegDiffusion.

## 1 Introduction

Gene regulatory networks (GRNs) describe the underlying regulatory network governing transcription and control of various cellular functions (Davidson and Levin, 2005; Karlebach and Shamir, 2008; Penfold and Wild, 2011). Understanding how these networks work can shed light on processes as diverse as organ development, response to inflammation, or oncogenesis. Moreover, understanding gene regulation from a systems perspective may help identify key points of regulation that could be amenable to therapeutic modulation (Emmert-Streib et al., 2014).

Despite increasing experimental data and a growing recognition of the binding sites of many regulatory proteins, GRNs are often inferred primarily from transcriptomic data (Mercatelli et al., 2020; Van Dam et al., 2018; Slonim, 2002). Such data sets offer unique functional *in vivo* snapshots of how genes are expressed in particular cells or under various conditions. Previously, such inference was based on microarray or bulk RNA sequencing data sets (Mercatelli et al., 2020; Schaffter et al., 2011; Hecker et al., 2009). More recently, the increasing accessibility of single-cell RNA sequence data has provided a higher resolution view of expression in particular cells and cell states (Nguyen et al., 2021). However, inferring GRNs from single-cell data is still challenging (Pratapa et al., 2020). In part, this is because GRN inference methods struggle with the dimensionality of the data. For data sets consisting of thousands of genes, there are potentially millions of edges to consider. Many algorithms do not scale well at this level. Noisy data further amplifies the challenge. Key among the recognized patterns of noise in single-cell data is “dropout” (Ghazanfar et al., 2016), where transcripts, disproportionately those with low or moderate expression levels, are lost from the expression counts. Other work has shown that even nonzero counts can be subject to technological background noise (Janssen et al., 2023).

Numerous methods, ranging from Bayesian Networks (BNs), mutual information (MI), to tree-based methods, have previously been proposed for GRN inference. BN-based methods, such as G1DBN (Lèbre, 2009) and ebdbNet (Rau et al., 2010), model GRNs as causal inference problems in a directed acyclic graph. MI-based methods, such as ARACNE (Margolin et al., 2006), CLR (Faith et al., 2007), MRNET (Meyer et al., 2007), and PIDC (Chan et al., 2017), measure the statistical dependence between two random variables and sort the edges according to these dependencies. Tree-based methods, such as GENIE3 (Huynh-Thu et al., 2010), dynGENIE3 (Huynh-Thu and Geurts, 2018), and GRNBoost2 (Moerman et al., 2019), rely on variable importance, a metric to rank variables while creating trees, to sort corresponding edges.

Recent advances in deep learning have offered new approaches to GRN inference. A recently published approach known as DeepSEM (Shu et al., 2021) learns the adjacency matrix by reconstructing the expression data through a modified Variational Auto-Encoder (VAE). The authors found that DeepSEM runs efficiently and outperforms many commonly used GRN inference methods on multiple benchmarks. Its network structure uses a parameterized adjacency matrix; an encoder, which transforms gene expression data to latent variables; and a decoder, which reconstructs the expression data from latent variables. The normality of the latent variables is enforced using the Kullback–Leibler (KL) divergence.

Yet, noise in single-cell data still presents challenges. Previously, we introduced the idea of dropout augmentation (DA) and presented a novel method called DAZZLE (Zhu and Slonim, 2023). The DA idea establishes a bridge between the “dropout” events in single-cell data and the deep learning regularization method also called “dropout” (Srivastava et al., 2014). Instead of trying to eliminate all spurious zeros in single-cell data, DAZZLE demonstrates the advantage of training the model with added zeros and explicitly predicting the (partially augmented) noise. Since a modest amount of augmented zeros can simulate dropout events, they increase model robustness and improve the benchmark performance of GRN inference. Encouraged by our positive findings, here we further explore the value of noise injection for GRN inference methods.

One emerging methodology that has shown potential in various domains, specifically computer vision, is that of diffusion probabilistic models (Ho et al., 2020; Sohl-Dickstein et al., 2015). By simulating the process of diffusion, or the spread of information, diffusion models are likelihood-based methods that aim to restore data from Gaussian corruption by an iterative process. In Denoising Diffusion Probabilistic Models (DDPMs) (Ho et al., 2020), a diffusion probabilistic model consists of a non-parameterized forward process, which adds small amounts of noise to the data for each of a number of steps, and a parameterized reverse process, which reconstructs a less-noisy version of the data. The reverse process is trained on predicting the added noise so that the de-noised data can be recovered by subtracting the predicted noise from the noisy data. Recent studies have suggested similarity between diffusion models and a generalized form of VAE with many latent spaces (Luo, 2022). While a classic VAE enforces normality in the center latent space with KL divergence, diffusion models enforce a trajectory to normality by the diffusion process. In addition, some studies further connect diffusion models with score matching models via annealed Langevin dynamics sampling (Song et al., 2021). This finding puts the diffusion process within the framework of stochastic differential equations (SDE) and enables continuous diffusion steps. To date, there are a few very recent preprints using diffusion models for single-cell data (Tang et al., 2023), and none of these consider modeling the regulatory relationships among genes.

Here, we introduce RegDiffusion, a novel diffusion probabilistic model focusing on the interactions among variables. To our knowledge, this is the first time a diffusion probabilistic model has been applied for this purpose. We demonstrate that GRNs can be learned effectively using the objective of predicting added noise in a diffusion process. The network architecture of our model is surprisingly simple, yet it outperforms several benchmark GRN inference methods. Overall, RegDiffusion has the following advantages:

- Compared to previous VAE-based models, the runtime of RegDiffusion improves from *O*(*m*^3^*n*) to *O*(*m*^2^), where *m* is the number of genes and *n* is the number of cells. One of the most important theoretical breakthroughs is the utilization of a more restrictive assumption allowing elimination of the costly adjacency matrix inversion step. Inference of networks with more than 15,000 genes now takes less than 5 minutes.
- RegDiffusion combines high-performance GRN inference with high stability. Compared to VAE-based deep learning models, RegDiffusion enforces a trajectory to normality by its diffusion process, which helps stabilize the learning process.
- Although presented as a deep learning method, RegDiffusion can be viewed as a form of Bayesian Network and is based on widely-used assumptions in gene regulatory research (Sanchez-Castillo et al., 2018).
- The success of RegDiffusion also illustrates the potential of applying diffusion models to single-cell and other kinds of noisy tabular data.

## 2 Methods

### 2.1 Problem Statement

Given a single-cell expression count table **X** ∈ ℝ^*n*×*m*^, where *n* is the number of cells and *m* is the number of genes, our objective is to infer the weighted adjacency matrix of the underlying gene regulatory network **A** ∈ ℝ^*m*×*m*^. It is reasonable to assume that the count table **X** reflects the typical zero-inflation and background noise affecting most single-cell expression technologies. We use **x** ∈ ℝ^*m*^ to denote the expression counts measured in one cell.

### 2.2 Denoising Diffusion Probabilistic Models

#### 2.2.1 Forward process

As outlined in the Introduction, a DDPM consists of a forward process and a reverse process. The forward process, denoted by *q*(**x**_*T*_|**x**_0_), is a non-parameterized Markov Chain that transforms an original unperturbed gene expression vector **x**_0_ to Gaussian noise **x**_*T*_ in *T* steps (eq. 1). This process generates a series of noisy samples **x**_0_, **x**_1_, …, **x**_*T*_ and as it proceeds, **x** gradually loses its distinct features as illustrated in Figure 1.

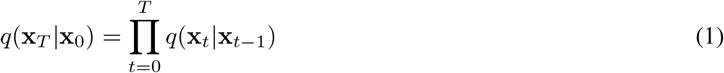

In each step of the forward process, *x*_*t*−1_ was perturbed according to a constant *β*_*t*_ ∈ (0, 1), which is part of a pre-generated noise schedule *β*. The value of *β*_*t*_ is usually very small. Intuitively, *β*_*t*_ represents the amount of information loss at each step. In the original DDPM paper (Ho et al., 2020), the authors proposed using the transformation function shown below (eq. 2; following their notation, the first parameter is simply the perturbed input at time *t*, while the second and third parameters are the mean and variance of the Normal distribution at that time). This transformation uses *β*_*t*_ to control both the mean and variance of the sampling process. The fact that 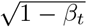 is always less than 1 ensures that after a number of steps, the mean of the final distribution will approach 0:

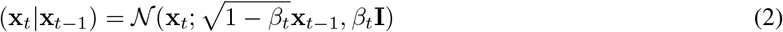

In practice, however, it is not feasible to do iterative sampling during training. To solve this problem, the authors of DDPM proposed a reparameterization trick that enables the generation of **x**_*t*_ from **x**_0_, or *q*(**x**_*t*_|**x**_0_), in one single step. We briefly explain this technique here, as it is crucial for our method.

**Figure 1:**
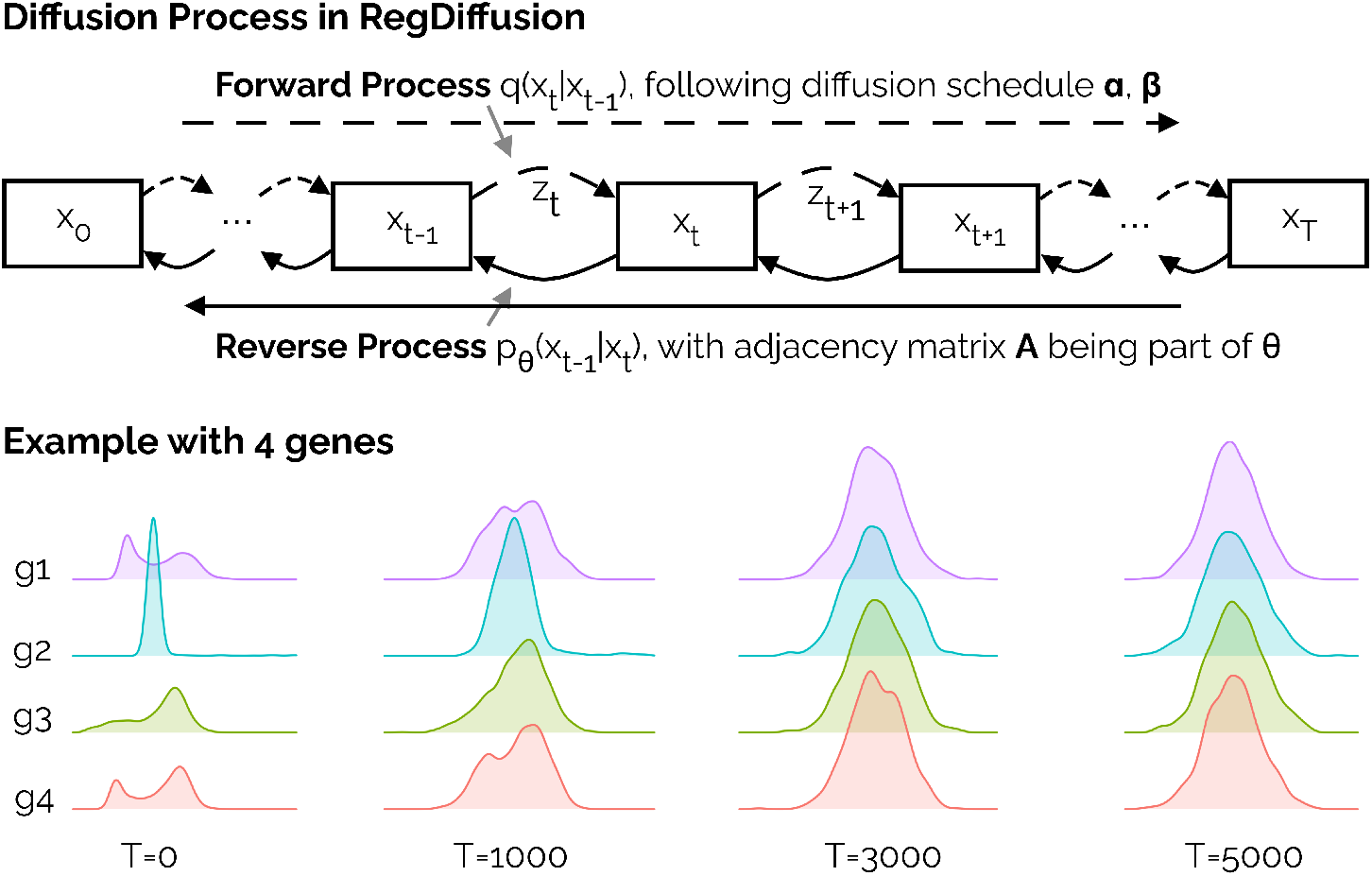
The Diffusion Process is an iterative process that consists of both the forward diffusion pass and the reverse pass. The top panel (based on (Ho et al., 2020)) shows the forward diffusion process transforming all data into Gaussian noise in T steps following a diffusion schedule, while the reverse process aims to recover the original distribution from the near-Normal distribution. We are modeling the reverse process with the adjacency matrix being part of the model. The bottom panel shows four representative genes and how their distributions change over time. Each gene begins with a unique distribution over all cells, but by the final time step the distribution is nearly Normal.

Let *α*_*t*_ = 1 − *β*_*t*_ and 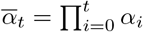. The reparameterization approach generates standard Gaussian noise **z** = 𝒩(0, 1). We can rewrite the transformation with the following equations:

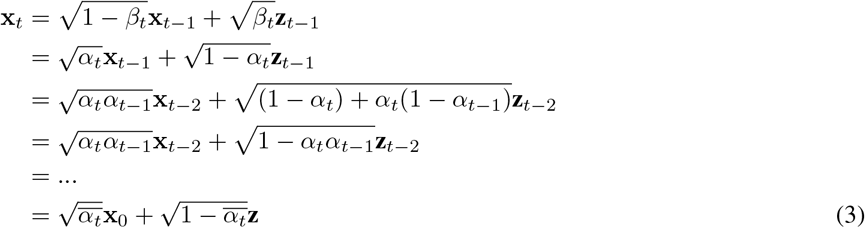

As mentioned above, the diffusion schedule *β* is pre-calculated. Therefore, the forward process at each iteration can be simplified as a non-parameterized function *f*_*forward*_ : **x**_**0**_ × **z** × *t* → **x**_**t**_. To be more specific, at each iteration, each cell gets a different time step *t*, which determines the strength of the perturbation. Then the forward process follows Equation 3 and generates the perturbed data **x**_*t*_ at the time step *t* using a randomly sampled standard Gaussian noise **z**.

#### 2.2.2 Reverse Process

Recall that the reverse process is a Markov Chain aiming to recover the original input **x**_0_ from **x**_*T*_. Since computing the exact reverse step *q*(**x**_*t*−1_|**x**_*t*_) is intractable, we use a parameterized model *p*_*θ*_(**x**_*t*−1_|**x**_*t*_) to approximate this process.

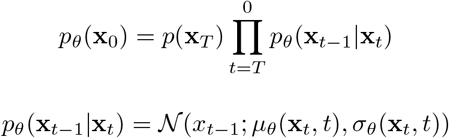

For simplicity, we chose to use the simplified loss function proposed in the DDPM paper:

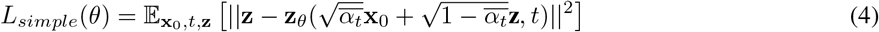

In other words, we need to create a model *p*_*θ*_(**z**|**x**_**t**_, *t*) to estimate the standard Gaussian noise **z** used in the forward pass (Eq. 3). In computer vision, this challenging problem is usually resolved using image segmentation models, such as U-Net (Ronneberger et al., 2015), to ‘segment’ the noise. Such an approach is not applicable in single cell data, where the data is tabular and the columns are unordered genes. Here, to resolve this problem, we propose a new way of calculating the noise using the linear additive assumption in Bayesian Networks.

### 2.3 RegDiffusion

#### 2.3.1 Graph noise estimating model

Our proposed RegDiffusion method relies on the linear additive assumption, which is commonly used in many GRN inference methods, especially in those based on Bayesian Networks (Friedman et al., 2000; Sanchez-Castillo et al., 2018). This assumption requires that the log probability of a particular gene is a linear combination of all of the regulator genes.

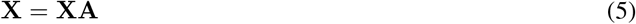

Note that this assumption is more restrictive than the one used in DeepSEM, which includes an addition term for random noise. The benefit of removing that term will become apparent later.

During the diffusion process, the gene expression matrix **X** is under slight Gaussian perturbation. The perturbed **XA** term on the RHS of Equation 5 will become a good approximation of the unperturbed **X** since the sum of the noise on all neighboring nodes will most likely trend back to zero with a much tighter variance.

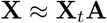

This means that the perturbed gene expression **X**_*t*_ can be written in the following form:

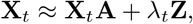

where **Z** is the Gaussian noise and *λ*_*t*_ is some constant for time step *t*. This equation can be easily transformed into a conceptual equation to estimate **Z**:

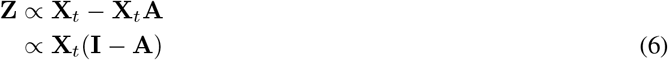

Equation 6 is exactly what we need to replace U-Net in the reverse modeling part of a diffusion model. This simple form describes a more straightforward network design compared with VAE-based models such as DeepSEM and DAZZLE. Most importantly, it is no longer necessary to perform matrix inversion on the (**I** − **A**) matrix. Matrix inversion, which is often solved by Gaussian elimination, runs in cubic time, and is one of the major bottlenecks in VAE-based models for large numbers of genes. At the same time, there is no need for separate encoders and decoders. All we need to do is to parameterize **X**_*t*_ and integrate the time step *t*.

The detailed structure of our proposed network, RegDiffusion, is shown in Figure 2. We explain each section in the following paragraphs.

**Figure 2:**
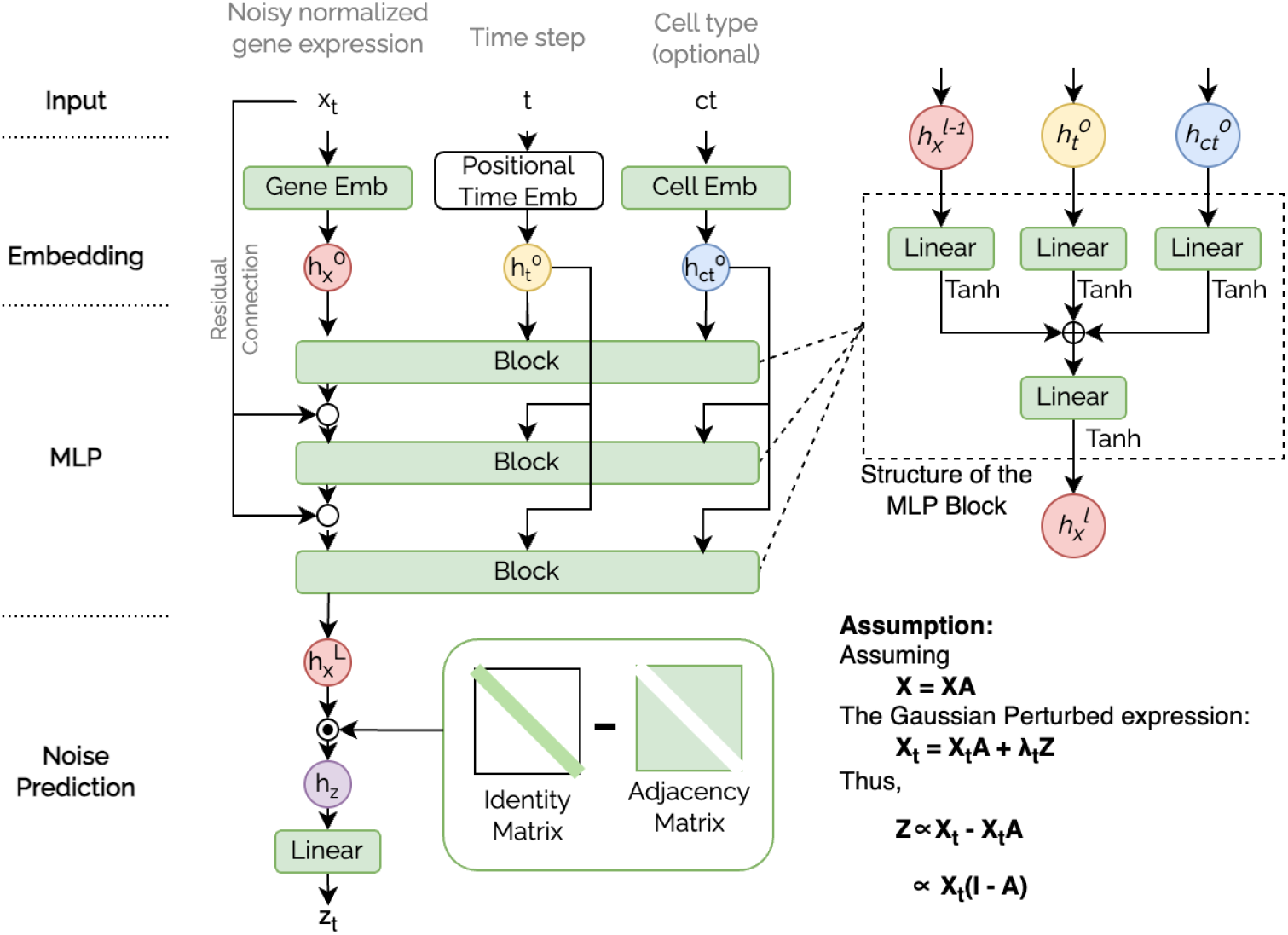
Graphic illustration of the RegDiffusion model. Green areas represent trainable parameters within the model. RegDiffusion starts with embedding the model inputs (gene expression *x*_*t*_, diffusion time step *t*, and cell type *ct*) into corresponding feature matrices (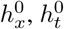, and 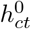). These feature vectors are then integrated through 3 layers of MLP training blocks. In each training block, the integrated gene features 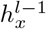 are mixed with the original embeddings of the time step and the cell type feature. The final integrated feature 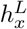 then multiplies with the (**I** − **A**) matrix and the outcome is used to predict the added noise *z*_*t*_.

#### 2.3.2 Data Preprocessing and Normalization

RegDiffusion requires a single-cell count table as input. We suggest users first follow common data preprocessing steps to address their specific needs. In experiments described here, we simply transform the data using the log2(count-plus-one) transformation and apply a standard quality assurance workflow (removing cells with high mitochondrial content or low/high UMI or gene counts). In terms of gene filtering, we removed genes that are not expressed at all. There is no need to restrict consideration to only the most variable genes, because RegDiffusion with all detected genes runs fairly quickly on modern GPUs.

We first normalize the expression table by cells using min-max normalization to balance inter-cell differences. Then, we z-score normalize the genes.

#### 2.3.3 Embedding

As shown in Figure 2, our model starts with an embedding process that transforms the input gene expression vector *x*_*t*_, diffusion time step *t*, and (optionally designated) cell type *ct* to corresponding embeddings 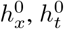 and 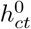. For each cell, the gene embedding 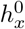 is a 2D array in the shape of 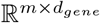, where *m* is the number of genes and *d*_*gene*_ is the size of the gene embedding. The first column is always the normalized expression values. The other columns are trainable gene features that are shared across cells. Next, following the standard practice for Diffusion Probabilistic Models, we use a Sinusoidal Positional Embedding to embed the time steps *t*. This process transforms an ordinal time step to a meaningful vector 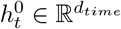 using sine and cosine transformation. Finally, RegDiffusion works well without additional cell type labels. However, if such information is available in existing data, users have the option to supply it to the model. In this case, the cell type embedding 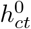 is anther 2D array in the shape of 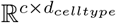, where *c* is the number of cell types and *d*_*celltype*_ is the dimension of the cell type embedding. Note that here, all the feature dimensions (*d*_*gene*_, *d*_*time*_, and *d*_*celltype*_) are hyperparameters of the model. In the current implementation, we set *d*_*gene*_ to be the same as the size of the first block.

#### 2.3.4 Feature Learning MLPs

In RegDiffusion, we use a set of multilayer perceptrons (MLPs) to aggregate feature matrices (*h*_*x*_, *h*_*t*_, and *h*_*ct*_) into a representation that is ready for noise prediction, as in Equation 6. To achieve this goal, we design an MLP building block as shown in Figure 2. This building block consumes the previous gene feature 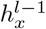, the original gene feature 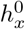, the time step embedding 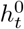, and the cell type embedding 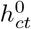. These features are converted into the same dimension using three separate linear models, and the activated values are summed together followed by another linear layer to build the new gene feature 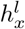. After *L* layers of the learning block, we are able to build a combined feature 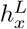.

#### 2.3.5 Adjacency Matrix Initialization and Regularization

The adjacency matrix in RegDiffusion is an *m* × *m* matrix of trainable model parameters, where *m* is the number of genes. Here we assume the average effect of a random gene to another, or the “regulation norm”, is 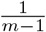. The initialization value of the adjacency matrix is set to five times the regulation norm to help the model explore both potential links and non-links. During the training process, unless otherwise specified, the learning rate on the adjacency matrix is set to one fiftieth of the regulation norm to ensure the learning steps on the adjacency matrix scale with the number of genes. To help the model converge faster, we also employ a soft threshold function (as shown in Equation 7) of half the regulation norm on the absolute values of the edge weights, so tiny edge weights are clipped during the training process. At the same time, the diagonal entries of the adjacency matrix are set to zeros to discourage self-regulation. To increase the sparsity of the adjacency matrix, we applied a combination of L1 and L2 losses on the matrix (weights controlled by two hyper parameters). To prevent overfitting and increase model robustness, we also apply dropout to the adjacency matrix; specifically, a proportion of edge weights are randomly set to zeros while the rest are scaled accordingly during training.

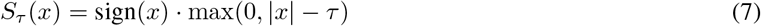

#### 2.3.6 Noise Prediction

Following equation 6, with the learned feature matrix *h*_*x*_ coming out of the stacked blocks and the adjacency matrix, we can calculate the feature of the added noise *h*_*z*_ using matrix multiplication. This feature is then converted into the prediction of noise *z* using a linear layer.

#### 2.3.7 Data Loading and Model Convergence

RegDiffusion considers data from each cell as an observation. In reality, the number of cells in different single-cell data varies significantly. For different datasets, if the model sees the entire dataset in every training iteration, it would take different numbers of iterations to converge. To overcome this issue, in each training iteration, we sample a fixed number of cells with replacement. This bootstrap resampling strategy also helps make RegDiffusion more robust.

The convergence point of RegDiffusion could be identified using the “amount of change” of the adjacency matrix with the help of the iteration training loss. The “amount of change” score is defined as the average change in value of the adjacency matrix after one iteration of training, in units of the regulation norm 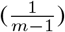. Eventually, this score will approach 0, reflecting that at this point in the training process, the optimizer is only making tiny updates on the adjacency matrix, showing that the model has converged.

As shown later in the result section, with a fixed sampling size per iteration, the convergence of RegDiffusion on different datasets follows a similar pattern, regardless of the number of genes and cells. The default number of training iteration is selected based on these observations.

#### 2.3.8 Network Interpretation and Neighbor/Neighborhood Extraction

After convergence, the next task is to interpret the inferred network. However, the difficulty of this task increases quadratically with the number of genes. Here, for the purpose of validating the inferred networks in their biological contexts, we propose visualizing the 2-hop local neighborhoods of specific interesting genes. For a given gene, we extract its top-k neighbors (sorted by the absolute values of their edge weights) as the first hop; then, for each gene in that set, we extract its top-k neighbors and add them to the “second hop” set. Finally, we scan through the neighbors of the second-hop genes, to add any remaining edges to nodes in either the first hop or second hop node sets. Thus, multiple genes that work together as a group are pulled closer in the visualization space, making their biological relationship more apparent.

### 2.4 Experiment Design

Experiments were run on the BEELINE benchmark data sets, where there are multiple, different putative “ground truth” networks to which to compare inferred regulatory edges. In addition, we inferred networks from two published sets of single-cell data in microglia, which can be compared to our knowledge about regulation in these cell types and which demonstrate the method’s feasibility in a more realistic context. Experimental details appear in the following subsections.

#### 2.4.1 Data Sets - BEELINE Benchmark

The BEELINE single-cell benchmark (Pratapa et al., 2020) comprises 7 distinct preprocessed single-cell RNAseq datasets from both human (hESC, hHep) and mouse (mDC, mESC, mHSC-E, mHSC-GM, mHSC-L), offering a broad scope for evaluation. With the provided benchmark-generation scripts in BEELINE, users can specify the number of target genes. BEELINE selects the most variable genes, based on the variance and p values returned by a fitted generalized additive model (GAM). BEELINE also suggests evaluating with respect to three sets of “ground truth” networks, namely the STRING v11 database (Szklarczyk et al., 2019), non-cell-type specific transcriptional networks, and cell-type specific networks. The non-cell-type specific network combines ChipSeq-based data collected from DoRothEA(Garcia-Alonso et al., 2019), RegNetwork(Liu et al., 2015), and TRRUST v2(Han et al., 2018), while the cell-type specific networks were experimentally generated by the BEELINE authors. In the main text, we use the STRING network as the ground truth and set the number of target genes to be 1,000. The exact numbers of genes/cells/true edges for each dataset are reported in Table 1. Our results on the other two ground-truth networks, yielding similar findings, are included in Supplement Tables 1-5. Since the cell-type-specific networks, though intuitively appealing, proved problematic for evaluation in several ways (see Section 3.4), we also examine cell specificity by comparing networks across data sets in different cells.

**Table 1:** RegDiffusion outperforms most GRN inference methods on BEELINE benchmark datasets (Metric: AUPRC Ratio; Ground truth: STRING; Number of target genes: 1000). Numbers reported are mean and std. of AUPRC Ratio compared with random guess over 10 runs. Higher ratios indicate better performance. Here, bold **Red** and italicized *orange* text indicate the best and *2nd best* algorithms. In this experiment, we ran each algorithm on all available data with the default settings.

|  | hESC | hHep | mDC | mESC | mHSC-E | mHSC-GM | mHSC-L |
| --- | --- | --- | --- | --- | --- | --- | --- |
| # of Genes | 1410 | 1448 | 1321 | 1620 | 1204 | 1132 | 692 |
| # of Cells | 758 | 425 | 383 | 421 | 1071 | 889 | 847 |
| # of True Edges | 5,149 | 9,000 | 5,898 | 8,479 | 1,826 | 1,311 | 154 |
| GENIE3 | 1.98 (0.01) | 1.86 (0.01) | **1.72 (0.01)** | 2.05 (0.01) | 4.19 (0.05) | 6.27 (0.04) | 7.02 (0.08) |
| GRNBoost2 | 1.67 (0.01) | 1.50 (0.01) | 1.43 (0.01) | 1.88 (0.02) | 3.65 (0.05) | 5.16 (0.11) | 7.07 (0.08) |
| DeepSEM* (single-run) | 1.95 (0.08) | 1.66 (0.10) | 1.57 (0.05) | 2.20 (0.03) | 5.00 (0.49) | 5.83 (0.81) | 6.69 (1.81) |
| DeepSEM (ensemble) | 2.11 (0.02) | 1.85 (0.02) | *1.67 (0.02)* | *2.31 (0.03)* | 5.67 (0.13) | *6.76 (0.17)* | *7.50 (0.12)* |
| DAZZLE | **2.50 (0.05)** | *1.87 (0.02)* | 1.60 (0.02) | 2.26 (0.06) | **6.02 (0.15)** | 6.39 (0.16) | **7.65 (0.12)** |
| RegDiffusion | *2.48 (0.01)* | **1.99 (0.01)** | 1.66 (0.01) | **2.45 (0.02)** | *5.96 (0.07)* | **7.14 (0.07)** | 7.28 (0.07) |

We compared the performance of RegDiffusion on these seven benchmark data sets to that of four other methods: GENIE3, GRNBoost2, DeepSEM, and our previously proposed method DAZZLE. GENIE3 and GRNBoost2 are two tree-based machine learning methods that have been found to perform well on single-cell data according to recent benchmarks (Kang et al., 2021; Pratapa et al., 2020). Particularly, as previously reported, with the same BEELINE benchmark we use, they are among the best three methods (together with PIDC (Chan et al., 2017)) and have been found to outperform SCODE (Matsumoto et al., 2017), SINCERITIES (Papili Gao et al., 2018) and PPCOR (Kim, 2015). DeepSEM and DAZZLE are both variational autoencoder based neural networks. For GENIE3 and GRNBoost2, we used the implementation from the Python package arboreto v0.1.6 (Moerman et al., 2019). For DeepSEM and DAZZLE, we used our implementation from the Python package GRN-dazzle v0.0.2 (Zhu and Slonim, 2023). Note that as described by the authors, DeepSEM is an ensemble algorithm that combines results from 10 runs. Because of its speed, we include the single-run version, denoted by DeepSEM*, in our comparisons as well as the ensemble version.

#### 2.4.2 Data Sets - Hammond Microglia Dataset

To assess RegDiffusion’s capacity on real-world single-cell data, we tested it on two different mouse brain microglia datasets. The first one is taken from a published single-cell data set on mouse microglia across the lifespan (Hammond et al., 2019) (Gene Expression Omnibus (Edgar et al., 2002) accession GSE121654). For our analysis, we used data from all four healthy male adults (P100) and filtered out cells with fewer than 400 or more than 3,000 unique genes, cells with more than 10,000 UMIs, and cells with over 3% of reads mapping to mitochondrial genes. For clarity and interpretability of results, we further removed mitochondrial genes, ribosomal genes, and pseudogenes with “Gm” prefixes. Finally, we removed genes that were not expressed at all in any of the cells. The final processed data consists of 8,258 cells and 14,065 genes. Note that this data set potentially includes many different microglia subtypes.

#### 2.4.3 Data Sets - Cerebellum Atlas Microglia Dataset

The second real-world microglia single-cell data is derived from a published transcriptomic atlas of mouse cerebellar cortex consisting of 611,034 cells of many types (Kozareva et al., 2021). The authors assigned cells to into known cell types based on specific gene markers in cell clusters. For microglia cells, the marker genes used were *Prkcd* and *C1qa*. From this dataset, we didn’t remove any cells from those designated as microglia, but we applied the same criteria as above to filter genes. We then log-plus-one transformed the count data. The final processed data set consists of 1,296 cells and 15,547 genes.

#### 2.4.4 Evaluation Metrics

Following the BEELINE paper, we use Area Under the Precision Recall Curve Ratio (AUPRCR) as the main evaluation metric. For the link prediction task as we have here, there is a huge class imbalance between the positive and negative group. As argued in (Davis and Goadrich, 2006), AUPRC is a better metric than Area Under the Receiver Operating Characteristic (AUROC) when dealing with class imbalance. AUPRC Ratio is simply the AUPRC score divided by that of a random predictor. Since the number of all possible edges and the number of true edges are usually very large in actual GRNs, the values of the AUPRC itself tend to be small. Converting them to ratios make it easier to understand the performance across benchmarks. We also report the ratio value for Early Precision (EP), which is essentially the top-k precision where k is the number of true edges. Results for the EP ratio are provided in Supplement Table 1, 3, and 5. Because it is a more familiar metric, the AUPRC ratio results are shown in the main text.

## 3 Experimental Results

### 3.1 RegDiffusion is Highly Accurate on BEELINE Benchmarks

Table 1 presents a performance comparison of AUPRC ratios for several GRN inference methods on the BEELINE benchmark data sets. The seven data sets differ in terms of the number of cells, genes, and edges in the ground truth networks - details appear in the table header.

The proposed RegDiffusion method generally outperforms most other GRN inference techniques across the different datasets, as indicated by higher AUPRC Ratios. It achieves top results for the hHep, mESC, and mHSC-GM data sets, and it is either the second or third best for the others, with performance above 95% of the top score in all cases.

Results for the EP Ratio metric also used in the DeepSEM paper appear in Supplement Table 1, 3, and 5. RegDiffusion performs even better by that metric, where it is either best or second best for all data sets. The Supplement contains results for different “ground truth” data sets and EPR or AUPRCR, but in all cases, RegDiffusion is either the clear winner, or one of the top performers along with DAZZLE and DeepSEM (ensemble).

Furthermore, the performance of RegDiffusion is very stable across runs, as indicated by the small standard deviations of AUPRC Ratios in 10 repeated runs. It has even better stability than the ensemble version of DeepSEM and is almost as stable as GENIE3 and GRNBoost2.

### 3.2 RegDiffusion Runs Faster than Competitors

As mentioned in Section 2, one of the most important breakthroughs in RegDiffusion is the elimination of the matrix inversion step on the adjacency matrix, which is required by previous VAE-based models. Matrix inversion requires cubic time and is the key computational bottleneck when the number of genes is large. By incorporating bootstrap sampling with a fixed pool size, the running time of RegDiffusion is independent of the number of cells. In section 3.5, we further show that this leads all models to converge in a fixed number of steps.

Compared to previous VAE-based models, the run time of RegDiffusion drops from *O*(*m*^3^*n*) to *O*(*m*^2^). A comparison of algorithm execution time is provided in Table 2. Across all benchmarks, RegDiffusion runs in a fraction of the time needed by any previous algorithm. This speedup has greater impact on larger data sets. For the two real-world single-cell data sets, each with tens of thousands of genes and thousands of cells, RegDiffusion takes only minutes, while previous VAE-based methods require hours of computation.

**Table 2:** Algorithm execution time (reported in seconds unless noted). Time cost on BEELINE data sets are averages over 10 runs. All code was executed on HPC computing nodes with 4 cores and 12GB memory. DeepSEM, DeepSEM*, DAZZLE, and RegDiffusion are executed with an additional NVIDIA A100 card.

|  | hESC | hHep | mDC | mESC | mHSC-E | mHSC-GM | mHSC-L | Hammond | Atlas |
| --- | --- | --- | --- | --- | --- | --- | --- | --- | --- |
| # of Genes | 1,410 | 1,448 | 1,321 | 1,620 | 1,204 | 1,132 | 692 | 14,065 | 15,547 |
| # of Cells | 758 | 425 | 383 | 421 | 1,071 | 889 | 847 | 8,258 | 1,296 |
| GENIE3 | 6,462 | 2,931 | 2,709 | 4,173 | 7,723 | 5,674 | 2,493 | - | - |
| GRNBoost2 | 2,055 | 992 | 962 | 2,171 | 2,442 | 1,382 | 379 | - | - |
| DeepSEM* (single run) | 29 | 15 | 12 | 17 | 30 | 23 | 15 | 4hr 21min | 57min |
| DeepSEM (ensemble) | 251 | 147 | 116 | 168 | 298 | 229 | 139 | 43hr 33min | 9hr 34min |
| DAZZLE | 31 | 16 | 13 | 18 | 34 | 27 | 19 | 4hr 47min | 1hr 10min |
| RegDiffusion | 7 | 8 | 7 | 9 | 6 | 5 | 5 | 3min 28s | 4min 10s |

### 3.3 Benchmark network interpretability example

While we lack space to analyze all the inferred networks on the benchmark data, here we examine an example: the network inferred from BEELINE’s mESC data (mouse embryonic stem cells) using the method described in section 2.3.8. For the choice of target gene, we picked *Hist1h1d* (or *H1*.*3*), a histone encoding transcription factor based on RegNetwork (Liu et al., 2015), as the target gene simply because it has the highest edge weight in the inferred adjacency matrix. Substantial expression and regulation of histone proteins in embryonic stem cells is unsurprising (Luger et al., 1997).

In Figure 3a, the inferred 2-hop neighborhood around Hist1h1d consists largely of two highly interconnected regions (“hairballs”, colored blue and yellow) with a layer of connecting genes. The blue histone complex includes 15 histone genes, comprising the entire set of histone genes available in the mESC dataset. RegDiffusion effectively retrieved all of them as direct neighbors using only expression data. The yellow cluster, which includes 5 minichromosome maintenance protein complex (MCM) genes among others, is enriched for links to to DNA replication. Separating the blue and yellow gene clusters is a layer of connecting genes, including *Rbbp4, Brca1, Kntc1, Dnmt1, Fbxo5*, and *Rad54b*. Among them, *Rbbp4* is a histone binding protein (Balboula et al., 2015). *Brca1* is a well-studied tumor suppressor gene that, in conjunction with histone h1, plays a crucial role in DNA repair (Ozgencil et al., 2023). The connection between *Dnmt1* and histone deacetylase has also been reported (Fuks et al., 2000). *Rad54b* deficiency has been reported to be associated with chromosome breakage (Russo et al., 2018).

**Figure 3:**
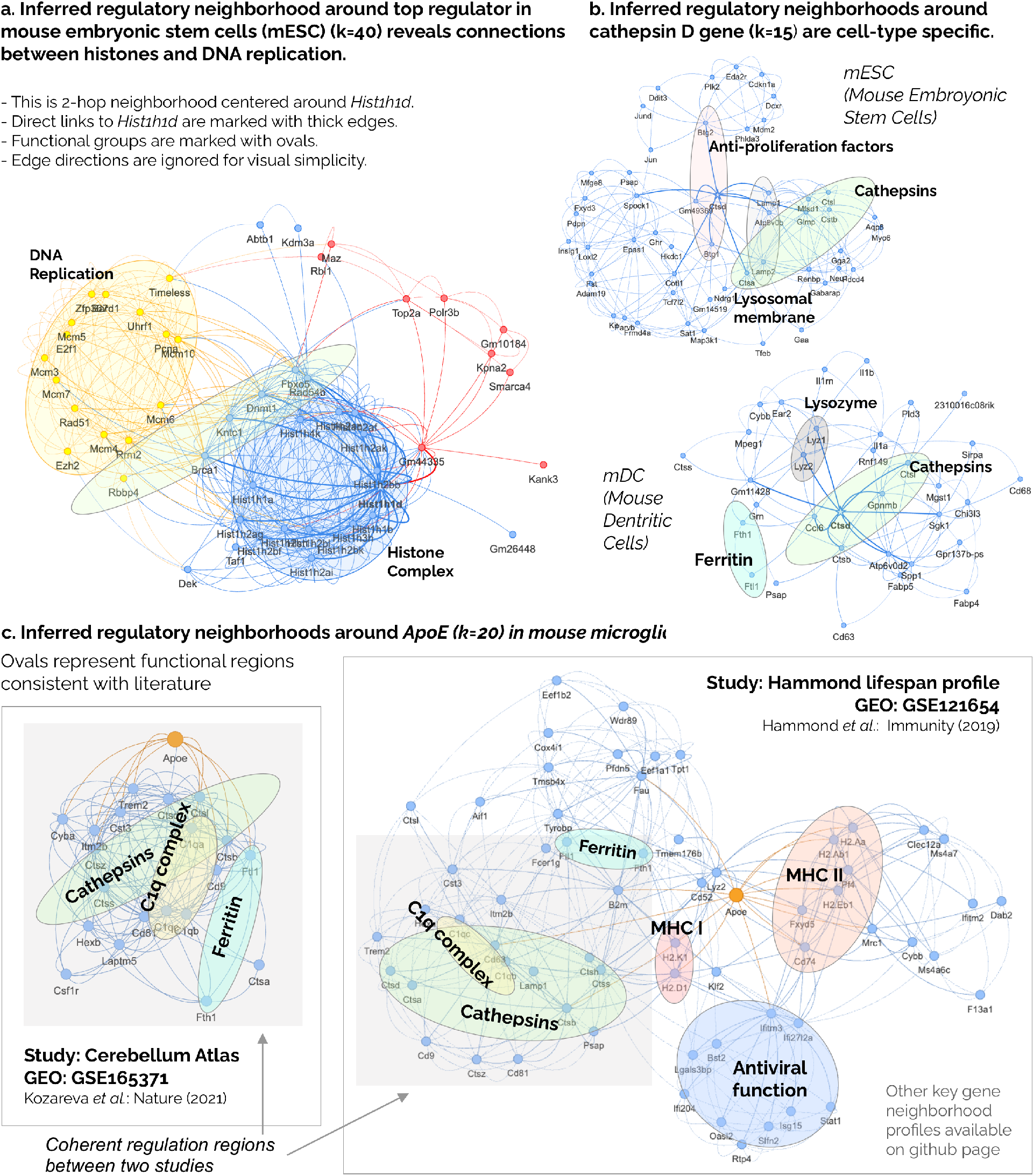
Local regulatory neighborhood analysis shows that the inferred regulatory networks generated by RegDiffusion are biologically interpretable. All the networks visualized here are 2-hop neighborhood around target genes at indicated top-k levels.

### 3.4 Inferred regulatory neighborhoods are cell-type specific

As mentioned in Section 2.4, we chose not to focus on the cell-specific ChIPSeq ground truth from BEELINE for a number of reasons. First, all of the unsupervised GRN inference methods we tested report performance near that of random guessing when compared to the cell-type specific data for the corresponding network. Second, although DeepSEM reports slightly better results than we found with most other methods (though still effectively no better than random guessing), that performance required a different set of hyperparameters than was otherwise suggested, one that minimizes the regularization on the adjacency matrix and minimizes the regulations of the KL divergence. The underlying cause of this pattern is still unclear, so we do not feel comfortable with the comparisons on these networks. Nonetheless, the comparable performance of RegDiffusion (under minimal adjacency matrix regularization) on this set of ground truth is reported in Supplement Table 4 and 5, to be interpreted with these caveats in mind.

To further address the challenge of cell-type-specific analysis, we compared the inferred regulatory neighborhoods around genes that are expressed across various cell types. For example, we noted that cathepsin genes are expressed in multiple datasets including mESC (mouse embryonic stem cells), mDC (mouse dendritic cells) from BEELINE, and both microglia datasets. As the most abundant lysosomal proteases, cathepsins play an important role in intracellular protein degradation and immune response (Yadati et al., 2020). As shown in Figure 3b, in the inferred networks in mESC, the top regulators of *Ctsd* include B Cell Translocation Gene (*Btg*) 1 and 2. *Btg1* and *Btg2* are anti-proliferation factors that regulate cell growth and differentiation. It is not surprising to see they trigger the expression of cathepsin to complete their function. In the inferred networks in mDC, one of the biggest changes is the disappearance of *Lamp1* and *Lamp2* from the neighborhood of *Ctsd* even though these two genes are expressed and measured in the mDC data. One possible explanation is that Lamp proteins in dendritic cells, or DC-LAMP, are biochemically different from the other *Lamp* proteins (Arruda et al., 2006; de Saint-Vis et al., 1998). Studies have shown that they are immediately fused into MHC II compartment and that this is a very cell-type-specific process for dendritic cells. In mouse microglia cells, as shown in Figure 3c, cathepsin genes (light green ovals) interact with many microglia homeostatic genes as part of the immune response. The RegDiffusion networks shown are consistent with these cell-specific changes.

Overall, the specificity of the inferred GRNs depends on the cell-type and cell state of the input data. When the input data has many cell types, the inferred GRN could be considered as overlaying many cell-specific GRNs but it also provides an overview of the cellular processes. With cell-specific data, the inferred GRNs might also be more specific, and there might be fewer pathways going through hub genes. We recommend that users design their project workflows based on their specific research questions.

### 3.5 Characteristics of RegDiffusion

The speedup obtained by RegDiffusion, in addition to relying on the avoidance of the matrix inversion step, comes from using bootstrapping to ensure convergence in time independent of the number of cells. In Figure 4a, by inspecting the amount of change in the adjacency matrix per iteration in the unit of the regulation norm, we show that across all of our experiments, whether on benchmark or real data sets, the training processes follow a similar pattern and the models converge after a fixed number of steps.

**Figure 4:**
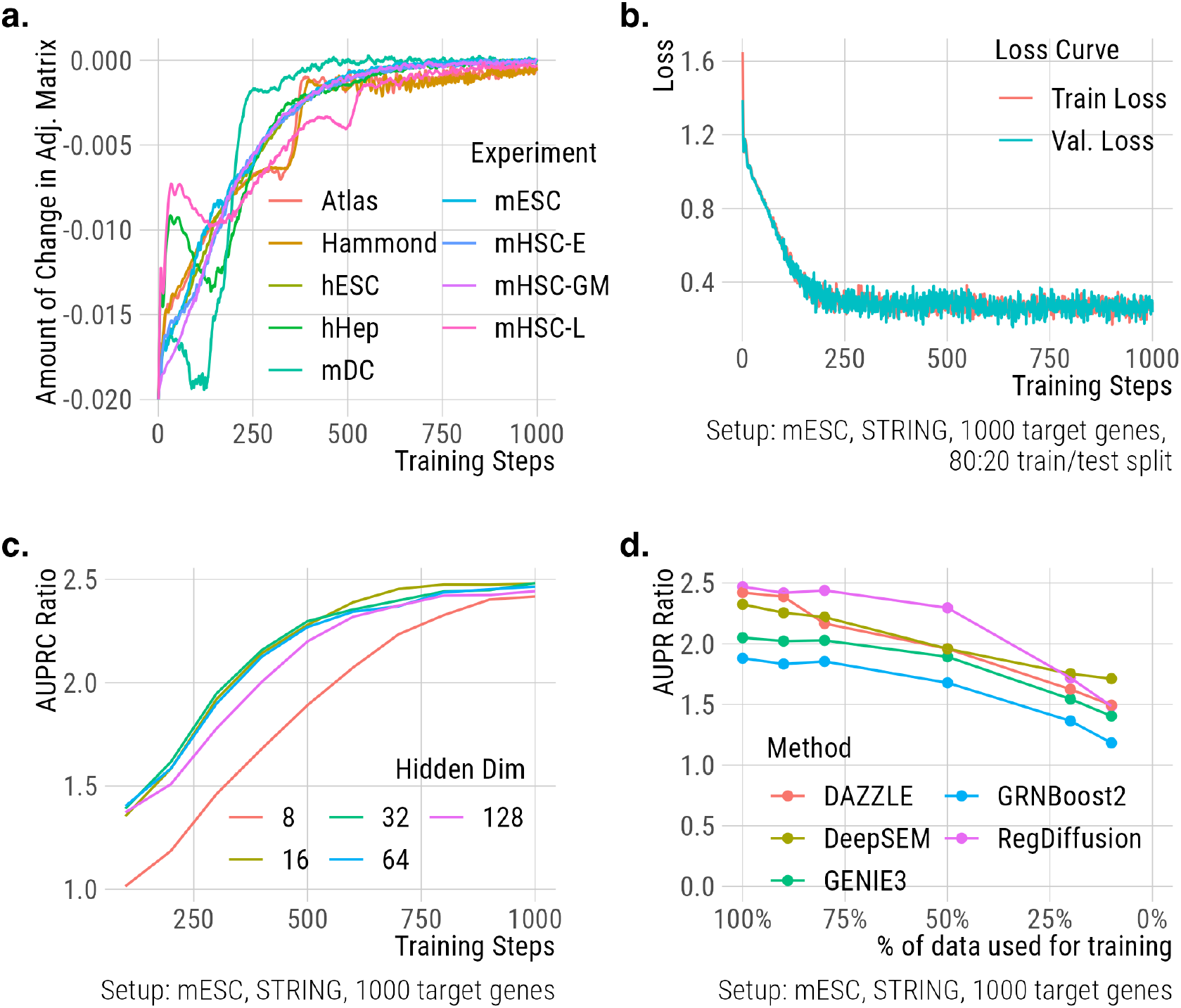
Properties of RegDiffusion. **a**. By measuring the amount of change in the adjacency matrix, we show RegDiffusion converges after a fixed number of steps in all experiments; **b**. Loss curves on train/test splits (4:1) show that the model was not overfitted. Also, it’s expected to see the training loss oscillating even after the model converges; **c**. Model performance is rather stable regardless of the hidden dimensions in the neural networks; **d**. Downsampling shows that RegDiffusion performs better than bulk RNAseq methods even when the sample size is limited.

Since RegDiffusion is trained only on the objective of predicting added noise, the ground truth GRN network never leaks into the training process. It works in an unsupervised manner in the same way as GENIE3 or GRNBoost2. To ensure there was no overfitting, we further investigated the training behavior by applying a train/test split on cells. Figure 4b illustrates the training losses of a typical training process on the mESC data set when we applied a 4:1 train/test split. As shown in the figure, the loss curves for the train and test splits are never separated too much, showing that the model was not overfitted. Also, note that since the diffusion steps are stochastic and it is usually harder to learn the fine details, it is expected to see training loss oscillating even after the model converges.

In terms of the number of parameters used in the model, we tested a number of choices for the dimension of the hidden layer in the MLP building block; results are shown in Figure 4c. For all dimensions greater than 8, the performance of the model was rather stable. Since we would like to reduce the number of parameters in the model as much as possible, we set the default model dimension to 16.

To assess RegDiffusion’s performance on networks with different sizes, we varied the number of target genes in BEELINE and assessed model performance for 100, 250, 500, 1000, and 2000 target genes. Overall, RegDiffusion appears to maintain a more stable performance across different numbers of target genes, as compared to DeepSEM. In most cases, training with more genes is helpful for RegDiffusion but it hurts DeepSEM’s performance in many cases. When the number of target genes is very limited (for example, 100), the performance of RegDiffusion seems to be slightly downgraded but is still better than the performance of DeepSEM in the same situation. A full comparison of RegDiffusion, DeepSEM, and GRNBoost2 on benchmarks of various sizes is provided in Supplement Figure 2.

Finally, we investigated the performance of RegDiffusion with limited numbers of cells. We gradually down-sampled the training data and then compared the AUPRC Ratios of results from RegDiffusion, GENIE3, and GRNBoost2. Figure 4d shows that even with 10% of the mESC data, around 42 cells, RegDiffusion still works thanks to the bootstrap resampling methods. It also outperforms tree-based methods, which are usually considered to be robust. In practice, some cell subtypes may only have a few sampled cells. This analysis shows RegDiffusion can be effective even in such cases. In Supplement Figure 1, we further include a visual comparison of the inferred local neighborhood from two different random batches of 90% cell samples. The results are very similar.

### 3.6 Regulatory Networks in Microglia

In both large microglia experiments, we systematically evaluated performance using both the full STRING v11 network and the combined non-cell-type-specific network provided in BEELINE as ground truth. In both experiments, RegDiffusion yields better AUPRC Ratio and Early Precision Ratio compared with DeepSEM according to both benchmarks. For example, in the Hammond experiment evaluated on the STRING network, the inferred network from RegDiffusion has an AUPRC ratio of 1.22 and Early Precision Ratio of 2.23 while the network from DeepSEM reports an AUPRC ratio of 1.17 and Early Precision Ratio of 1.68. The full results are provided in Supplement Table 6.

In addition to benchmark testing, we further examined the biological interpretability of the inferred GRNs in mouse microglia using data from the two complementary studies described in Section 2.4. For each study, we inferred a GRN and analyzed the local regulation neighborhood around particular genes. For this discussion, we focus on *ApoE* (apolipoprotein E) and visualize the gene’s RegDiffusion-inferred network neighborhoods from both studies’ data in Figure 3c. *ApoE* is a well studied lipid regulator that plays a role in several diseases and disorders, but it is probably best known for the E4 allele’s association with Alzheimer’s Disease (AD) (Fernández-Calle et al., 2022).

The ApoE neighborhood inferred from the Hammond dataset is essentially a superset of the neighborhood inferred from the Cerebellum Atlas data. This is possibly because the microglia from the Atlas dataset were selected from clusters marked by expression of marker genes *C1qa* and *Prkcd*. Some microglial subtypes may have been filtered out in this process, or there might have been fewer microglial cells because of the focus on cerebellum and the large number of other cell types considered in the atlas.

Nonetheless, many key relationships are shared across the data sets in Figure 3c, and many inferred links have previously been experimentally confirmed. For example, the regulatory relationship between *ApoE* and MHC (major histocompatibility complex) type I genes, such as *H2*.*K1* and *H2*.*D1* has been recently confirmed in (Zalocusky et al., 2021) via an ApoE perturbation experiment. Activated Response Microglia (ARMs), a microglia subtype discovered in (Frigerio et al., 2019), have been found to have overexpressed MHC type II genes, including *H2-Aa, H2-Ab1*, and *Cd74*. RegDiffusion even further identified *H2*.*Eb1* as another potential biomarker for this cell type. On the right side of the MHC II complex in Figure 3c, we have another pair of genes *Mrc1* and *Ms4a7*. These two genes were identified in (Hammond et al., 2019) as biomarkers of a unique microglial state during the embryonic phase. While the data we analyzed is from adult mice, the topological position of these two groups suggests a possible relationship between the ARMs cell type and that embryonic cell state. We further identified connections between *ApoE* and a set of antiviral response genes, including *Ifitm3*, which connects directly to *ApoE* based on our model. *Ifitm3* has previously been identified as a *γ*-secretase modulatory protein that increases the production of amyloid-*β* in AD ((Hur et al., 2020)).

If we compare the inferred *ApoE* neighborhoods between the two studies, we can identify coherent shared patterns. These recapitulate previously identified links among *ApoE* and several functional gene groups, including the *C1q* (Complement component 1q) complex (*C1qa, C1qb*, and *C1qc* (Habenicht et al., 2022*)), Cathepsins (such as Ctsd, Ctsb*, and *Ctss* (Zhou et al., 2006), *(Samokhin et al*., *2010), (Li et al*., *2024)), Ferritins (Ftl1* and *fth1* (Ayton et al., 2015)), *and several microglia homeostatic genes (such as Trem2* (Wolfe et al., 2018), *Itm2b*(Biundo et al., 2015), *and Hexb*. While we didn’t find prior evidence explicitly showing ApoE regulating *β*-hexosaminidase (*Hexa, Hexb*), there is evidence that they are both implicated in, and have correlated expression in, AD pathology (Tiribuzi et al., 2011; Sierksma et al., 2020), suggesting such a relationship is plausible. Further, *β*-hexosaminidase plays a role in lysosomal function and autophagy. Our recent work examining fetal microglia in the setting of maternal immune activation(Batorsky et al., 2024) shows that the CLEAR (Coordinated Lysosomal Expression and Regulation) pathway, particularly *Hexa, Hexb*, and several cathepsins, is differentially regulated in microglia exposed to inflammation. From the consistency and plausibility of these networks, we conclude that the results from running RegDiffusion on current single-cell data sets are reasonable and consistent.

For comparison purposes, the local networks inferred by DeepSEM are provided in Supplement Figure 3. Overall, DeepSEM yields similar findings on the Hammond dataset. However, the MHC I group and the antiviral group are missing. There are more microglia homeostatic genes but there are also more orphan links that may be false predictions. For the Atlas dataset, DeepSEM generates a lot of orphan links but it still captures the *C1q* complex and the cathepsins among the noise. In addition to the local neighborhood around *ApoE*, we also provide visualizations for 10 other microglia signature genes on our project website.

## 4 Conclusion

We have presented RegDiffusion, a novel diffusion probabilistic model designed to infer GRNs from single-cell RNAseq data. RegDiffusion simulates the information diffusion process by iteratively adding Gaussian noise following a fixed schedule. Based on the benchmarking results, compared to other top methods, RegDiffusion demonstrates consistent superiority in GRN inference capacity while remain stable across runs. It also achieves a significant runtime reduction over comparable methods. Even with limited data, RegDiffusion still outperforms competitors. At the same time, by visualizing a local gene neighborhood in the inferred GRNs, we illustrate that GRNs inferred by RegDiffusion are consistent with current knowledge of molecular function.

Following the idea of Dropout Augmentation (Zhu and Slonim, 2023), which simulates the dropout phenomenon in single cell data by inserting zero values into expression data using Bernoulli sampling, RegDiffusion is our second attempt at leveraging noise injection in single cell analysis. The success of RegDiffusion in GRN inference also suggests several potential future research paths. First, it would be very interesting to explore the effects of RegDiffusion and its learned embeddings on other single-cell analysis tasks. Another important future research direction is understanding the inferred networks. Our proposed visualization method works well for inspecting individual genes, but it would be nice to investigate ways to understand the graphs more systematically. Finally, it would be helpful to find a solution to learn detailed features in single-cell data given the existence of background noise. In our experiments, we observed that the models are good at learning crude features (at high diffusion steps) but still have some difficulty learning fine features at low diffusion steps, a phenomenon that has been observed many times in other diffusion models.

Diffusion models in general have sparked a lot of research interest in recent years thanks to their huge success in computer vision. Together with a few recent pre-prints in 2023 (Tang et al., 2023), RegDiffusion presents as one of the initial attempts to solve the problems in single-cell data with diffusion models and is the first one that considers gene regulatory relationships. We anticipate that RegDiffusion many not only offer a reliable and convenient tool for GRN inference, but may also advance our understanding of regulatory genomics and single-cell analysis.

## 5 Supplement document for RegDiffusion

### 5.1 Alternative Benchmark Results

**Table S1:** BEELINE Benchmarking Results (**STRING network**, 1,000 target genes) evaluated on Early Precision Ratio (**EPR**). Results are from the same experiment shown in Table 1 in the main text.

|  | hESC | hHep | mDC | mESC | mHSC-E | mHSC-GM | mHSC-L |
| --- | --- | --- | --- | --- | --- | --- | --- |
| # of Genes | 1410 | 1448 | 1321 | 1620 | 1204 | 1132 | 692 |
| # of Cells | 758 | 425 | 383 | 421 | 1071 | 889 | 847 |
| # of True Edges | 5,149 | 9,000 | 5,898 | 8,479 | 1,826 | 1,311 | 154 |
| GENIE3 | 3.66 (0.07) | 3.17 (0.03) | 2.19 (0.04) | 3.48 (0.03) | 7.00 (0.06) | 8.53 (0.09) | 7.34 (0.14) |
| GRNBoost2 | 3.46 (0.09) | 2.78 (0.04) | 2.08 (0.04) | 3.69 (0.05) | 6.21 (0.06) | 7.32 (0.12) | 7.12 (0.24) |
| DeepSEM* | 4.12 (0.17) | 3.00 (0.23) | 2.13 (0.16) | 3.89 (0.13) | 7.27 (0.36) | 7.79 (0.65) | 6.92 (1.11) |
| DeepSEM | 4.55 (0.08) | 3.69 (0.05) | 2.16 (0.07) | 4.12 (0.07) | 7.87 (0.17) | 8.73 (0.24) | 7.48 (0.14) |
| DAZZLE | 5.31 (0.12) | 3.50 (0.13) | 2.04 (0.06) | 3.89 (0.09) | 7.99 (0.13) | 8.39 (0.12) | 7.57 (0.18) |
| RegDiffusion | 5.38 (0.05) | 3.51 (0.05) | 2.23 (0.04) | 4.18 (0.04) | 8.41 (0.09) | 9.19 (0.07) | 7.51 (0.30) |

**Table S2:** BEELINE Benchmarking Results (**Non-celltype-specific ChIP-Seq** network, 1,000 target genes) evaluated on AUPRC Ratio (**AUPRR**). Results are from a similar experiment to that shown in Table 1 in the main text, but with a different ground truth network.

|  | hESC | hHep | mDC | mESC | mHSC-E | mHSC-GM | mHSC-L |
| --- | --- | --- | --- | --- | --- | --- | --- |
| # of Genes | 1410 | 1448 | 1321 | 1620 | 1204 | 1132 | 692 |
| # of Cells | 758 | 425 | 383 | 421 | 1071 | 889 | 847 |
| # of True Edges | 4,597 | 5,335 | 3,918 | 8,030 | 1,960 | 1,358 | 317 |
| GENIE3 | 0.97 (0.00) | 1.01 (0.01) | 1.56 (0.01) | 1.65 (0.01) | 2.54 (0.02) | 3.56 (0.03) | 2.78 (0.04) |
| GRNBoost2 | 1.01 (0.01) | 1.09 (0.01) | 1.36 (0.02) | 1.49 (0.02) | 2.39 (0.04) | 2.84 (0.03) | 2.45 (0.05) |
| DeepSEM* | 1.20 (0.06) | 1.35 (0.07) | 1.71 (0.06) | 1.57 (0.05) | 2.95 (0.19) | 3.05 (0.24) | 3.06 (0.51) |
| DeepSEM | 1.24 (0.01) | 1.41 (0.01) | 1.94 (0.03) | 1.65 (0.02) | 3.18 (0.03) | 3.52 (0.14) | 3.15 (0.08) |
| DAZZLE | 1.27 (0.01) | 1.42 (0.02) | 1.80 (0.05) | 1.64 (0.03) | 3.39 (0.06) | 3.52 (0.16) | 3.32 (0.11) |
| RegDiffusion | 1.18 (0.01) | 1.38 (0.01) | 1.82 (0.02) | 1.72 (0.01) | 3.17 (0.03) | 3.69 (0.05) | 3.02 (0.08) |

**Table S3:**
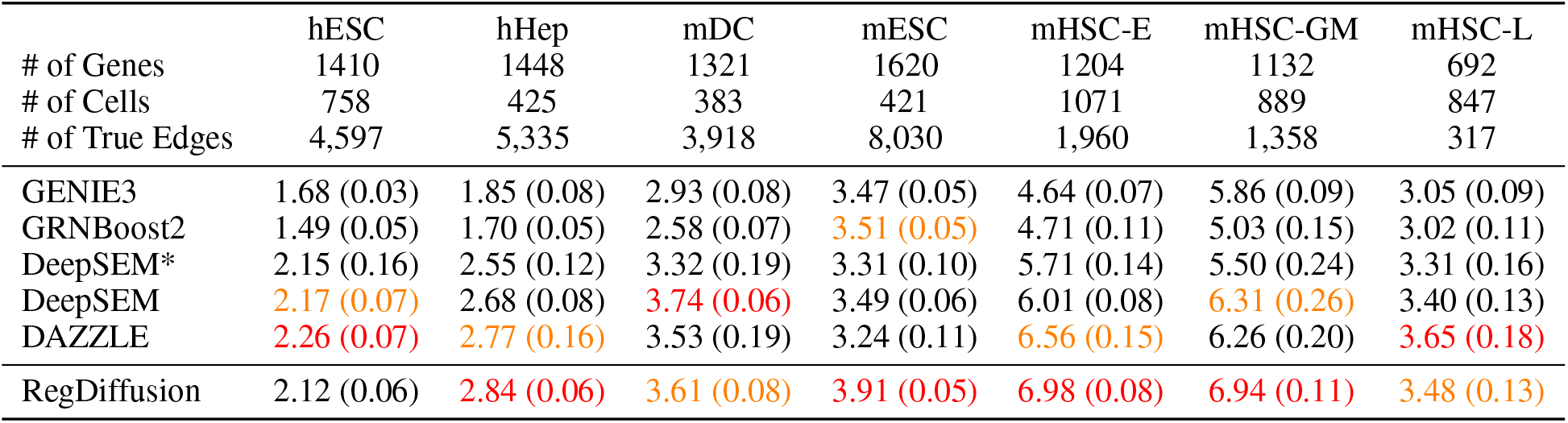
BEELINE Benchmarking Results (**Non-celltype-specific ChIP-Seq** network, 1,000 target genes) evaluated on Early Precision Ratio (**EPR**). Results are from a similar experiment to that shown in Table 1 in the main text, with a different ground truth network.

**Table S4:** BEELINE Benchmarking Results (**Celltype-specific ChIP-Seq** network, 1,000 target genes) evaluated on AUPRC Ratio (**AUPRR)**. DeepSEM*, DeepSEM, DAZZLE are all run using a different set of hyperparameters recommended by the DeepSEM paper only for use on this benchmark (L1 Loss = 1% of default; KL divergence = 1% of default). RegDiffusion also uses a different set of hyperparameters (L1 loss = 1% of default; L2 Loss = 1% of default; Adjacency matrix initialization = 10 * regulation norm). Because of the low performance ratios and unique hyperparameters here, plus a lack of understanding for why performance with the original hyperparameters is much worse, we do not recommend evaluating on this set of ground truth.

|  | hESC | hHep | mDC | mESC | mHSC-E | mHSC-GM | mHSC-L |
| --- | --- | --- | --- | --- | --- | --- | --- |
| # of Genes | 1410 | 1448 | 1321 | 1620 | 1204 | 1132 | 692 |
| # of Cells | 758 | 425 | 383 | 421 | 1071 | 889 | 847 |
| # of True Edges | 7,050 | 15,410 | 1,193 | 42,795 | 21,975 | 14,135 | 5,180 |
| GENIE3 | 0.96 (0.00) | 1.10 (0.00) | 1.01 (0.01) | 1.06 (0.00) | 0.99 (0.00) | 0.98 (0.00) | 1.05 (0.00) |
| GRNBoost2 | 1.01 (0.00) | 1.02 (0.00) | 1.00 (0.01) | 1.02 (0.00) | 0.99 (0.00) | 0.99 (0.00) | 1.01 (0.00) |
| DeepSEM* | 1.37 (0.05) | 1.13 (0.01) | 0.99 (0.07) | 1.03 (0.01) | 1.00 (0.01) | 0.98 (0.01) | 1.09 (0.02) |
| DeepSEM | 1.38 (0.01) | 1.14 (0.01) | 1.02 (0.02) | 1.03 (0.00) | 1.00 (0.00) | 0.98 (0.00) | 1.08 (0.01) |
| DAZZLE | 1.30 (0.12) | 1.13 (0.03) | 1.03 (0.09) | 1.02 (0.01) | 1.02 (0.01) | 0.99 (0.06) | 1.10 (0.03) |
| RegDiffusion | 1.37 (0.02) | 1.15 (0.01) | 1.04 (0.01) | 1.05 (0.00) | 1.02 (0.00) | 0.95 (0.01) | 1.08 (0.01) |

**Table S5:** BBEELINE Benchmarking Results (**Celltype-specific ChIP-Seq** network, 1,000 target genes) evaluated on Early Precision Ratio (**EPR**). DeepSEM*, DeepSEM, DAZZLE are all run using a different set of hyperparameters recommended by the DeepSEM paper only for use on this benchmark (L1 Loss = 1% of default; KL divergence = 1% of default). RegDiffusion also uses a different set of hyperparameters (L1 loss = 1% of default; L2 Loss = 1% of default; Adjacency matrix initialization = 10 * regulation norm). Because of the low performance ratios and unique hyperparameters here, plus a lack of understanding for why performance with the original hyperparameters is much worse, we do not recommend evaluating on this set of ground truth.

|  | hESC | hHep | mDC | mESC | mHSC-E | mHSC-GM | mHSC-L |
| --- | --- | --- | --- | --- | --- | --- | --- |
| # of Genes | 1410 | 1448 | 1321 | 1620 | 1204 | 1132 | 692 |
| # of Cells | 758 | 425 | 383 | 421 | 1071 | 889 | 847 |
| # of True Edges | 7,050 | 15,410 | 1,193 | 42,795 | 21,975 | 14,135 | 5,180 |
| GENIE3 | 0.95 (0.01) | 1.12 (0.01) | 0.99 (0.06) | 1.06 (0.00) | 1.02 (0.00) | 1.03 (0.09) | 1.07 (0.00) |
| GRNBoost2 | 0.78 (0.02) | 0.27 (0.00) | 0.97 (0.08) | 0.33 (0.00) | 0.21 (0.00) | 0.23 (0.01) | 0.44 (0.01) |
| DeepSEM* | 1.62 (0.05) | 1.12 (0.02) | 0.97 (0.19) | 1.01 (0.01) | 1.05 (0.00) | 1.06 (0.01) | 1.07 (0.01) |
| DeepSEM | 1.64 (0.01) | 1.12 (0.01) | 1.02 (0.10) | 1.01 (0.00) | 1.05 (0.00) | 1.07 (0.00) | 1.07 (0.00) |
| DAZZLE | 1.51 (0.17) | 1.12 (0.03) | 1.04 (0.15) | 1.02 (0.02) | 1.04 (0.01) | 1.02 (0.06) | 1.08 (0.03) |
| RegDiffusion | 1.65 (0.03) | 1.14 (0.01) | 1.10 (0.05) | 1.04 (0.00) | 1.02 (0.01) | 1.03 (0.01) | 1.05 (0.01) |

### 5.2 Training with Limited Data

#### 5.2.1 Result stability when training with limited number of cells

In addition to Figure 4d in the main text, we assessed the stability of the inferred networks when RegDiffusion is trained with a limited number of cells.

#### 5.2.2 Training with limited number of genes

Here we vary the number of target genes in BEELINE benchmark datasets and assess the inference capacity of RegDiffusion when training with limited or abundant data. Overall, RegDiffusion yields better results when more genes are available. When the number of target genes are limited, the performance of RegDiffusion gets impaired but overall, it’s still doing better compared to DeepSEM.

**Table S6:**
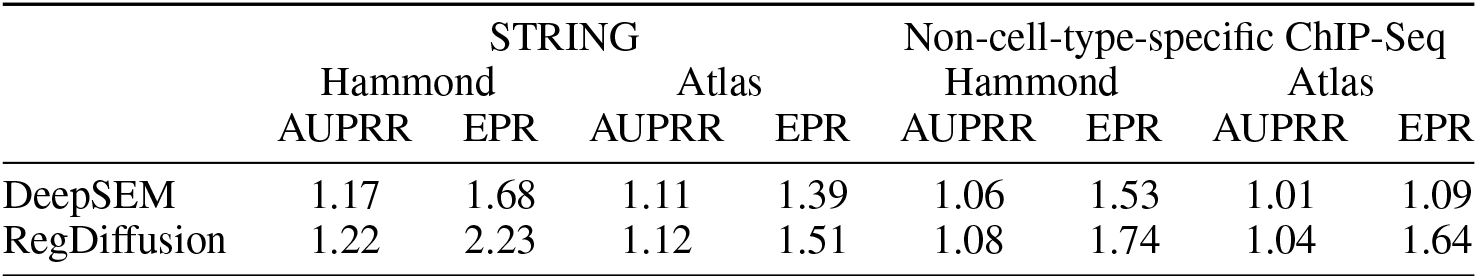
RegDiffusion outperforms DeepSEM in two large microglia experiments on all metrics. Both of the ground truth networks (STRING and Non-cell-type-specific ChIP-Seq are the full networks from BEELINE.

|  | STRING |  |  |  | Non-cell-type-specific ChIP-Seq |  |  |  |
| --- | --- | --- | --- | --- | --- | --- | --- | --- |
|  | Hammond<br>AUPRR | EPR | Atlas<br>AUPRR | EPR | Hammond<br>AUPRR | EPR | Atlas<br>AUPRR | EPR |
| DeepSEM | 1.17 | 1.68 | 1.11 | 1.39 | 1.06 | 1.53 | 1.01 | 1.09 |
| RegDiffusion | 1.22 | 2.23 | 1.12 | 1.51 | 1.08 | 1.74 | 1.04 | 1.64 |

**Figure S1:**
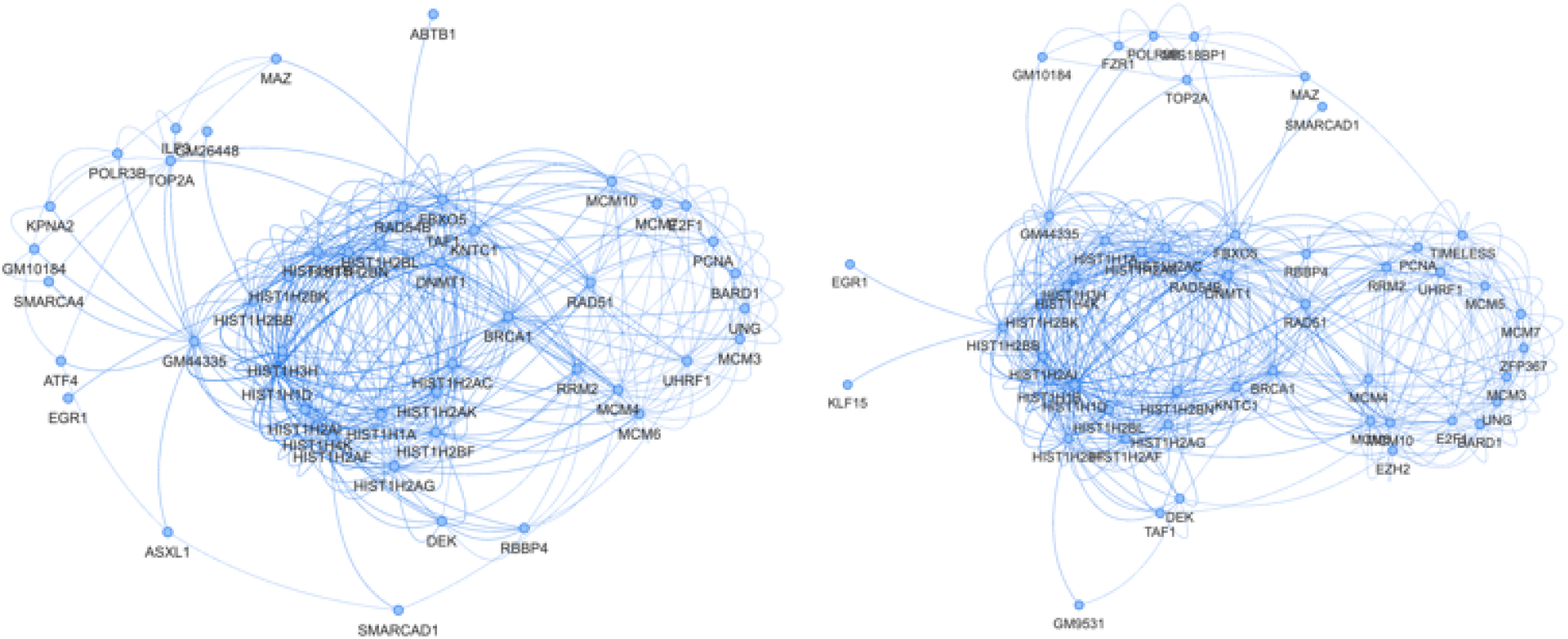
Inferred Local Neighborhood around Hist1h1d when inferred with two different batches of 90% sampled cells. These two neighborhoods are very similar with only some small variances (84-86% overlapping nodes and 65-71% overlapping edges).

### 5.3 Inferred Microglial Network using DeepSEM

**Figure S2:**
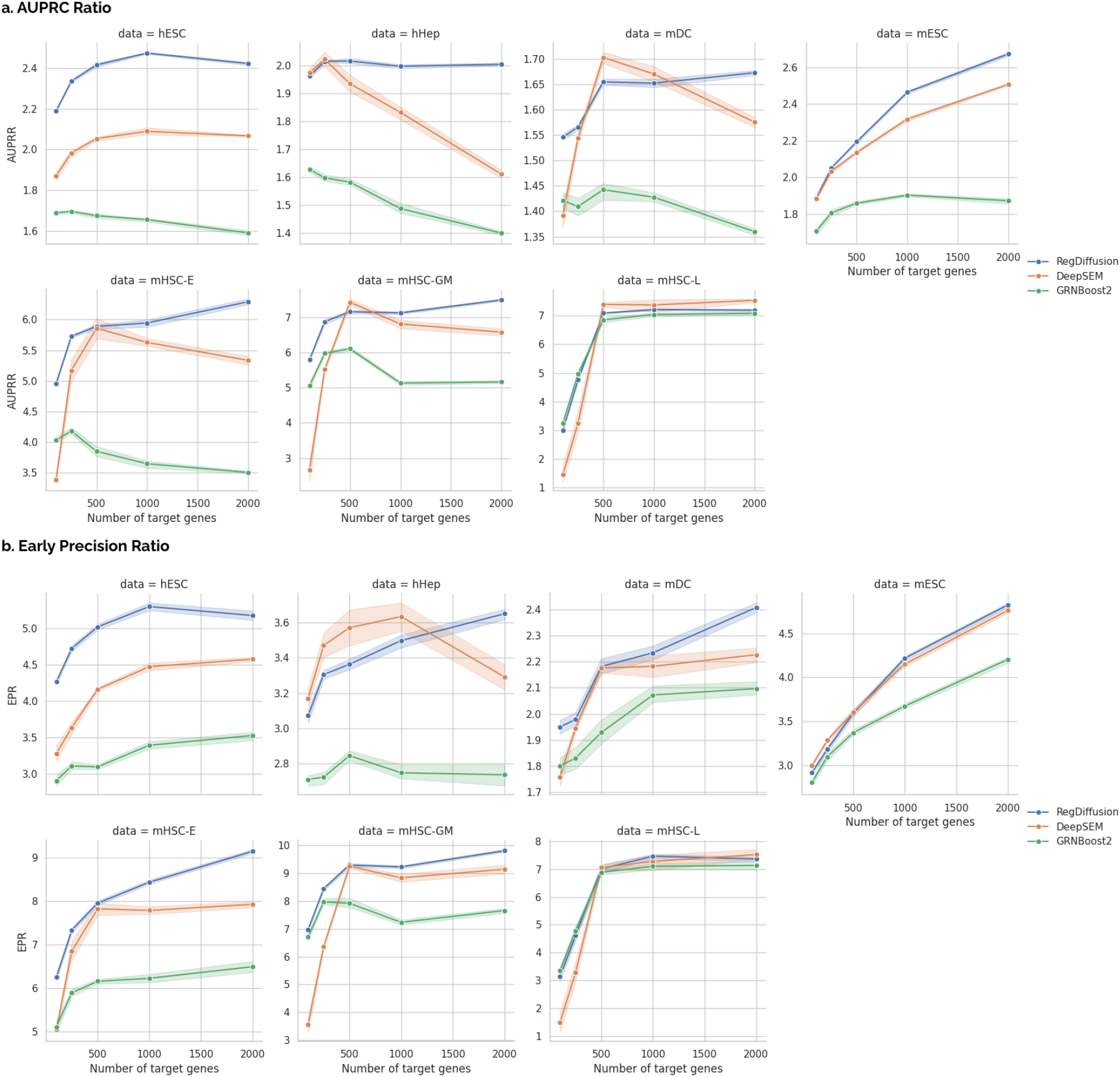
Comparison of Inference Performance (a. AUPRC Ratio; b. Early Precision Ratio) with Limited or Abundant data. Ground truth network is STRING. Line and bands represent mean and standard deviations of 10 runs. Each run of DeepSEM ensembles 10 single runs.

**Figure S3:**
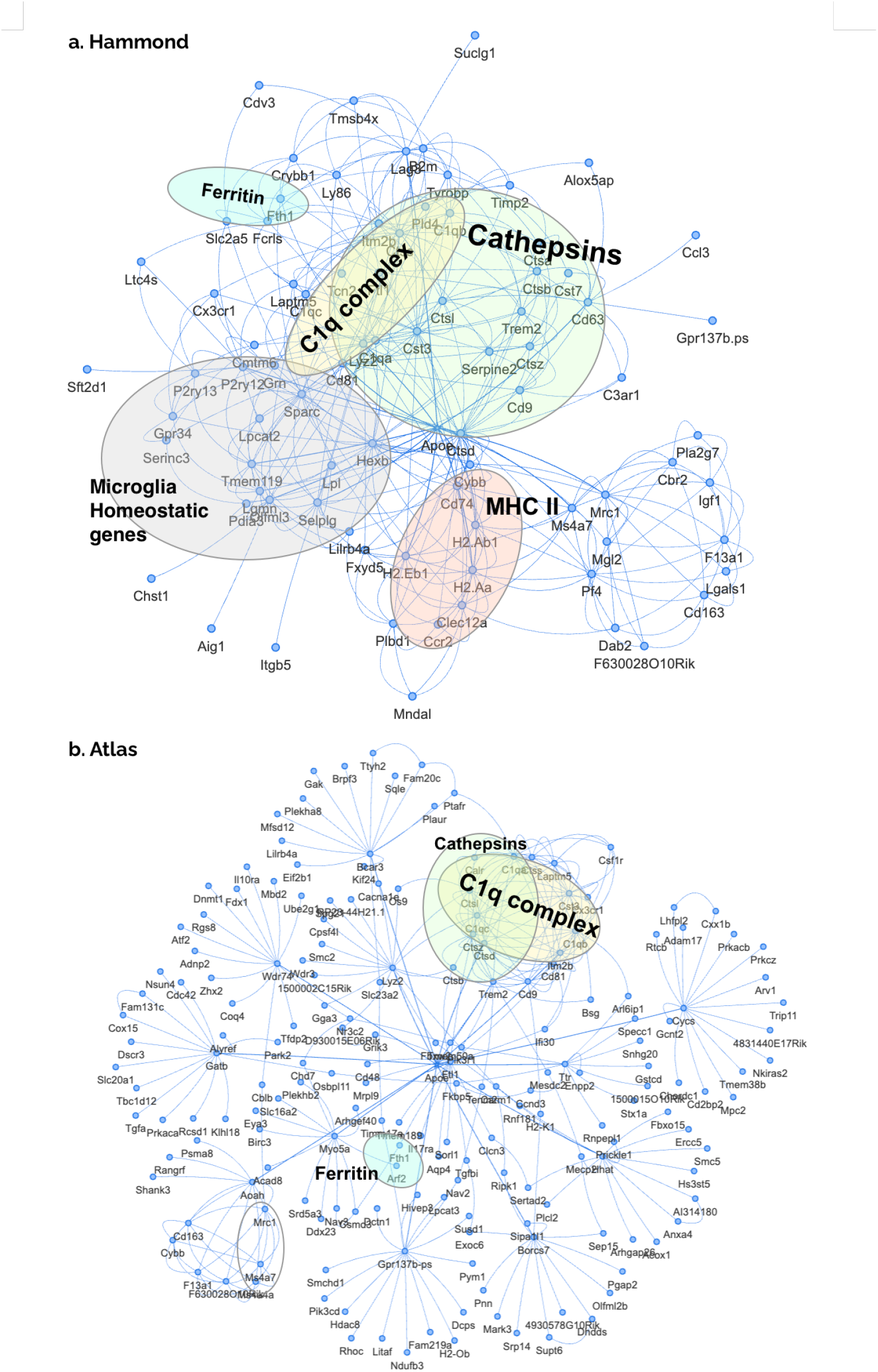
Inferred 2-Hop Regulatory Networks (k=20) around *ApoE* from (a. Hammond; b. Cerebellum Atlas) microglia data using DeepSEM. Overall, these two networks are similar to the ones inferred by RegDiffusion but in both cases, there are more orphan links that are probably false predictions. In addition, in the Hammond result, the MHC I group and the antiviral group are missing.

